# Scarf: A toolkit for memory efficient analysis of large-scale single-cell genomics data

**DOI:** 10.1101/2021.05.02.441899

**Authors:** Parashar Dhapola, Johan Rodhe, Rasmus Olofzon, Thomas Bonald, Eva Erlandsson, Shamit Soneji, Göran Karlsson

## Abstract

The increasing capacity to perform large-scale single-cell genomic experiments continues to outpace the computational requirements to efficiently handle growing datasets. Herein we present Scarf, a modularly designed Python package that seamlessly interoperates with other single-cell toolkits and allows for memory-efficient single-cell analysis of millions of cells on a laptop or low-cost devices like single board computers. We demonstrate Scarf’s memory and compute-time efficiency by applying it to the largest existing single-cell RNA-Seq and ATAC-Seq datasets. Scarf wraps memory-efficient implementations of a graph-based t-stochastic neighbour embedding and hierarchical clustering algorithm. Moreover, Scarf performs accurate reference-anchored mapping of datasets while maintaining memory efficiency. By implementing a novel data downsampling algorithm, Scarf additionally can generate representative sampling of cells from a given dataset wherein rare cell populations and lineage differentiation trajectories are conserved. Together, Scarf provides a framework wherein any researcher can perform advanced processing, downsampling, reanalysis, and integration of atlas-scale datasets on standard laptop computers.

## INTRODUCTION

The rapid evolution, integration, and diversification of high-throughput single-cell genomic technology continue to have a critical impact on our conceptual understanding regarding tissue heterogeneity, cell type specification, and differentiation ^1–3^. Accumulating technological advances reach beyond gene expression quantification to the measurement of higher-dimensional genomic features and an exponential increase of input cell numbers ^4^. This ability to generate large quantities of single-cell genomic data has in turn necessitated the development of new methods to facilitate appropriate data handling, processing, visualization, and analysis ^5^. Simultaneously, the rapid accumulation of datasets calls for possibilities to integrate results from different technological or biological origins. However, Sc-genomic data from large-scale studies are often made available through web portals where quick but limited access is granted and hence, limiting their full potential. Additionally, massive “atlas-level” datasets ^6^ demand to be analysed on servers with extensive memory capacity, as even loading the data becomes a bottleneck using desktop computers.

Thus, a major objective for computational analysis of Sc-genomic data has been to improve the scalability of analysis for increasing cell numbers and features ^7^. While the focus has been to improve computation time, memory scalability has largely been ignored even though memory capacity in computing systems is currently the major hurdle for increased usage of Sc-genomic datasets. Larger datasets with more than 100,000 cells obligate the use of specialized hardware with larger primary memory capacity (random access memory, RAM) in the order of several hundreds of gigabytes ^8^. This resource barrier prevents a majority of biologists and bioinformaticians hassle-free access to their own as well as publicly available datasets.

To meet the computing challenges associated with large datasets we have developed Scarf, a highly memory-efficient software solution allowing for analysis of single-cell genomic data in computing systems with low memory (RAM) capacity, such as standard laptop computers. Scarf is a modularly developed and extensible Python package, designed to work with any kind of single-cell genomic dataset presented as a matrix of cells and features. By creating a novel graph-based method for data down-sampling (downsampling henceforth), we have leveraged Scarf’s memory efficiency while preserving archetypes or differentiation trajectories. Scarf performs major steps of data analysis like data filtering, normalization, feature selection, linear dimension reduction, cell-cell neighbourhood graph creation, non-linear dimension reduction, clustering, and identification of discriminatory features. In the current version, functionality to analyse ScRNA-Seq ^9^, ScATAC-Seq ^10,11^, and CITE-Seq ^12^ datasets are included.

Efficient and interpretable clustering of cells is crucial for single-cell genomic analysis. To enable clustering at low memory consumption, a hierarchical graph clustering algorithm, Paris ^13^, scalable to millions of cells is introduced in Scarf. For visualization of generated cell clusters, the Uniform Manifold Approximation and Projection (UMAP) ^14^ algorithm is complemented with a graph t-distributed stochastic neighbour embedding (t-SNE) ^15^. When the same initialization of data is used the UMAP will highlight the magnitude of cluster relations while t-SNE will reveal the heterogeneity within the data.

Experimental perturbation of single cells can give answers to many research questions as well as elucidating cell responses in disease. Dissecting these contrasting conditions is not trivial as such datasets often generate distinct separate clusters when merged. Scarf is equipped with features for optimal data integration where samples are projected onto each other cell by cell, using a K-Nearest neighbour (KNN) mapping approach. This strategy avoids the generation of non-biological sample-separation when datasets generated from different technologies or techniques need to be combined, or from strong molecular signals when comparing cells after for example perturbation experiments or malignant transformation.

Importantly, benchmarking performed utilizing the largest publicly available scRNA-Seq and scATAC-Seq datasets show that Scarf can handle up to four millions of cells and up to a million of features on a regular laptop in a time-efficient manner while producing results that are consistent with the widely used existing, memory-exhausting tools.

## RESULTS

### Scarf enables analysis of even the largest scRNA-Seq and scATAC-Seq datasets on laptop computers

Single-cell genomic datasets undergo two stages of processing. In the first stage, sequencing reads are used to generate a count matrix which serves as the foundation for all the downstream analysis. The two axes of the count matrix usually consist of cell barcodes and features. For example, in single-cell RNA-Seq data, features normally represent genes/transcripts and for single-cell ATAC-Seq data features are the peak coordinates. Scarf achieves memory efficiency by dividing a dataset in small chunks and then compressing these chunks individually and storing them onto the disk (figure 1A). Chunks with a larger number of zero values automatically achieve a larger degree of compression. In contrast to other single-cell libraries like Loompy and Scanpy ^16^, Scarf uses the Zarr file format ^17^ to store chunked datasets and not HDF5 ^18^. Zarr improves support and performance for parallel operations as well as interactions with datasets that are stored remotely (see Methods). Learning algorithms like PCA (principal component analysis) and KNN (k-nearest neighbours) usually require the whole dataset as input. Scarf uses out-of-core (a.k.a. incremental) implementations of these algorithms that allow iterative input of data in small chunks. These incremental learning algorithms lead to the creation of the cell-cell neighbourhood graph structure of the data (see Methods for details) which can subsequently be used for downstream steps like generating UMAP/t-SNE visualizations, pseudotime ordering, etc.

**Figure 1:**
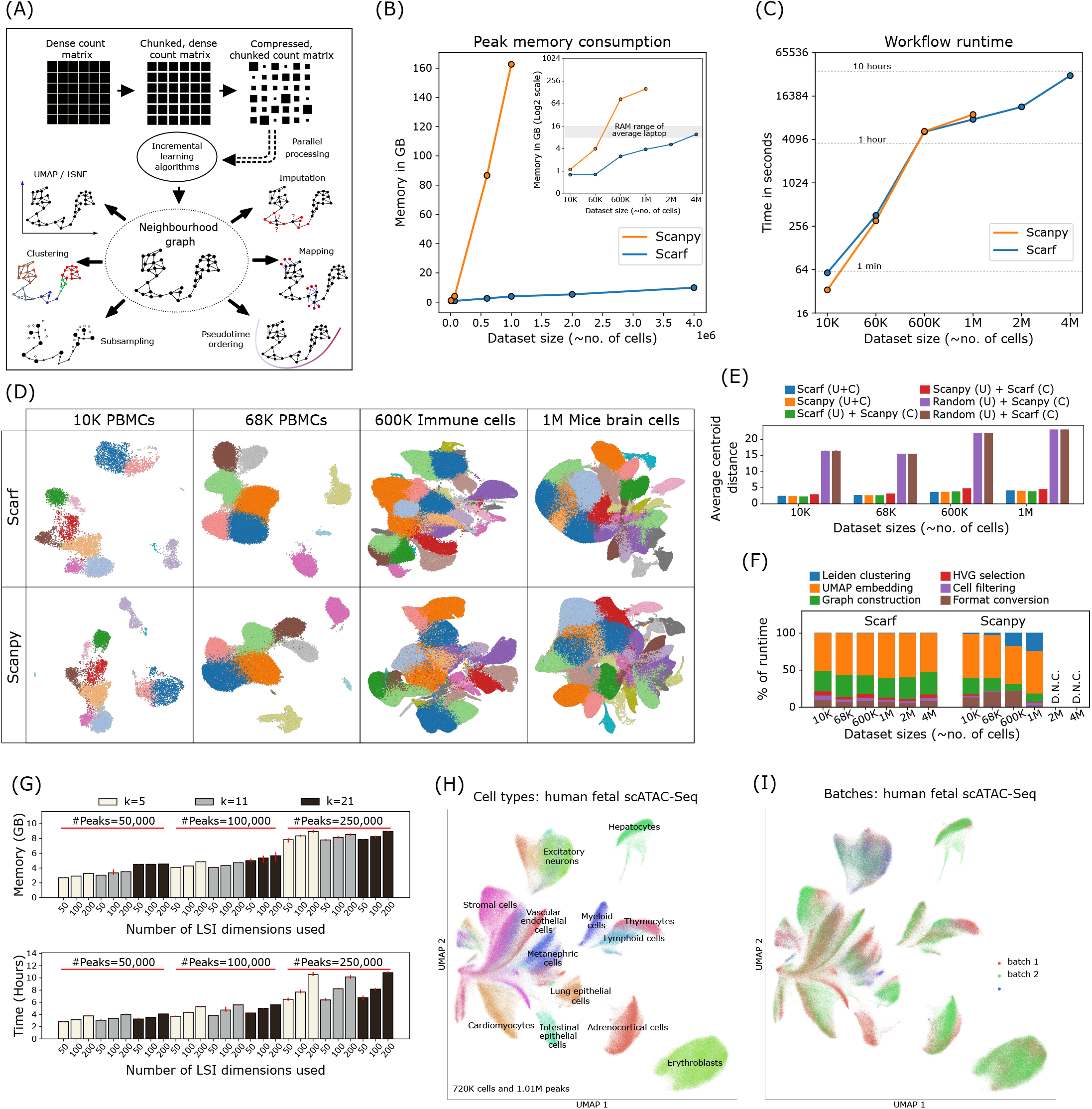
Scarf performs memory and time efficient computation to produce consistent embedding and clustering. **(A)** Schematic of the workflow of Scarf wherein the input data is illustrated as a matrix used to generate a cell-cell neighbourhood graph. Outward pointing arrows from the neighbourhood graph indicate the operations that can be performed on the graph in no particular order. **(B)** Plot showing the amount of memory consumed by Scarf and Scanpy on datasets containing up to 4 million cells. The inset image shows the same data with y-axis on log2 scale. Dots connecting the lines indicate the number of cells on the x-axis and corresponding memory consumed on the y-axis. Lines are drawn to indicate general trend. **(C)** Plot showing the amount of time (in seconds) consumed by Scarf and Scanpy on the six datasets used for benchmarking. The x-axis shows the number of cells in the datasets as categorical labels. Horizontal dotted lines indicate the time consumed (in hours). **(D)** Plots showing UMAP embedding of cells calculated using Scarf and Scanpy. Cells are coloured, for both Scarf and Scanpy, by the cluster identity obtained using Scarf’s Leiden clustering. Only four of the six datasets, that were successfully processed using Scanpy are shown here. **(E)** Bar plots showing the average distance (in UMAP space) of cells from their corresponding cluster centroids. ‘U’ = UMAP and ‘C’ = clustering. **(F)** Percentage of time consumed by six broad steps in the processing pipeline of Scarf and Scanpy. D.N.C = ‘did not complete’ due to out-of-memory error. Please note that the Leiden clustering step might not be visible for Scarf when zoomed out because of its quick runtime. **(G)** Bar plots showing the memory and time consumption of Scarf on the >700K single-cell ATAC-Seq dataset. Each bar shows results of benchmarking conducted using a combination of multiple number of peaks, k (number of neighbours in the nearest neighbour graph) and number of latent sematic indexing dimension used over which the graph was computed. The error bars show the standard deviation computed over three iterations of the entire pipeline. **(H)** Scatter plot showing UMAP embedding of the cells from the >700K single-cell ATAC-Seq dataset. Cells are coloured by author-annotated cell types or **(I)** batches of data generation.

One of the most common objectives of a single-cell analysis pipeline is to generate a low dimensional embedding of cells (e.g., UMAP and t-SNE) including partitioning of cells into distinct clusters. Along with UMAP/t-SNE embedding and clustering, current protocols involve common processes like data normalization, feature selection, dimension reduction, and cell neighbourhood graph generation, forming a basic workflow of single-cell data ^19^. We benchmarked the time and memory consumption of such a basic workflow using Scarf and another widely used scRNA-Seq analysis toolkit in Python, Scanpy ^16^.

Using six scRNA-Seq datasets ^20–22^ with increasing cell numbers, we found that under the same analysis parameters (see Methods), Scarf had substantially lower memory consumption than Scanpy (figure 1B). Importantly, using less than 16GB of memory (RAM), which is commonly available in modern laptops, even the largest set of four million cells ^22^ were efficiently processed by Scarf. In contrast, Scanpy was not able to process the two and four million cell datasets due to high memory consumption, despite 200GB of RAM being available during the benchmarking experiments. For the largest set successfully analysed by Scanpy (one million brain cell dataset generated by 10x genomics), approximately 40 times more computational memory was used compared to Scarf, under the same parameters and equivalent steps. Moreover, the lower memory consumption by Scarf did not come at a cost of the runtime of the workflow, which was similar to Scanpy (figure 1C). Of note, Scarf was able the process all analysis steps using the four million cell dataset within 10 hours without exceeding 16GB of memory usage.

The four datasets that could be processed by Scanpy, were visualized by the UMAP embeddings generated by either Scarf or Scanpy (figure 1D). To aid the visual inspection, cells were coloured based on their cluster identity obtained using Scarf. The generated UMAPs indicate that very similar embeddings of cells were achieved. To quantify any potential differences, we calculated the average centroid distance (ACD), i.e., the average Euclidean distance of cells from their cluster centroids in the UMAP space, and cross-evaluation was performed using UMAP from one and clustering information from the other pipeline. Indeed, ACD values were found to be similar when comparing the two pipelines to the ones obtained when both UMAP and clustering information was obtained from the same pipeline (figure 1E). In contrast, the ACD values were substantially higher after generating a random embedding of cells.

Next, extensive benchmarking was conducted under different combinations of three variable parameters: number of highly variable genes (HVGs), number of PCA components and neighbours to be used in the construction of cell-cell neighbourhood graphs. We found that memory consumption was consistently substantially lower in Scarf than Scanpy, across all the parameters tested (figure S1A-B). The number of HVGs and PCA dimensions used had a very low impact on memory consumption and runtime. However, using larger numbers of neighbours for graph building (parameter ‘k’) led to a substantial increment in memory consumption. We investigated the runtime of each major stage of the pipeline and found that the proportion of time consumed by each stage was similar across the benchmarked datasets (figure 1F). With the chosen parameters, UMAP was the longest running step for both Scanpy and Scarf, occupying more than 50% of the runtime. For Scanpy, the runtime for Leiden clustering was augmented when dataset size was increased, while consistently low in Scarf.

Scarf can easily be applied to other single-cell genomics methods, including large-scale scATAC-Seq. To demonstrate this scalability, we used a dataset generated from human foetal samples representing 15 organs, 720 613 cells in total and a feature set comprising 1.05 million peak regions ^23^. Scarf was able to process this data, generating clusters and UMAP embeddings, in approximately 5.5 hours using less than 2.5GB of RAM. As done previously for the scRNA-Seq datasets, we benchmarked the runtime and memory consumption across three critical parameters: number of peaks, latent semantic indexing (LSI) components used, and number of neighbours identified for neighbourhood graph creation (figure 1G). In contrast to RNA-Seq, we found that when the number of peaks and/or LSI components was increased, memory usage was increased. However, the memory consumption stayed under 10GB even when approx. 25% of peaks were included in the analysis. UMAP embeddings labelled with author-determined cell types (figure 1H) revealed that the UMAP embedding generated by Scarf accurately captured the heterogeneity of the cell types and categorized cells based on tissue of origin. Furthermore, the UMAP embedding, and the clustering of cells were not affected by batches of data generation (figure 1I).

### Topology assisted cell downsampling using Scarf

The increasing generation of high-throughput single-cell genomic data has prompted the use of downsampling (sub-selection of representative cells) to allow for advanced downstream analysis. Algorithms that have a polynomial space complexity (aka memory complexity) greater than *O*(*n*^2^) are generally not scalable to atlas scale datasets. Downsampling is used to obtain biologically informative results from high space complexity algorithm to either reduce the runtime or memory consumption ^24^. Random sampling is often used as a downsampling solution however it provides no guarantees of preserving archetypes or differentiation trajectories in the downsampled versions of the data. Recently, GeoSketch ^24^ has been proposed to solve this challenge, however, GeoSketch is not memory efficient as it has been designed to be used with large-scale computational resources. Here, a novel downsampling algorithm called TopACeDo (Topology Assisted Cell Downsampling) is embedded into Scarf. TopACeDo leverages the neighbourhood graph structure of the cells to perform downsampling that aims to conserve rare cell types, reduce the proportion of highly represented cell types, and preserve the underlying manifold of the transcriptional space. Briefly, TopACeDo identifies landmark points in the graph, seed cells, and then tries to find paths to connect those seed cells using prize-collecting Steiner tree algorithm (PCST) (figure 2A). An implementation of PCST with near-linear time complexity ^25^ is used to achieve fast and scalable downsampling on millions of cells (see Methods for details).

**Figure 2:**
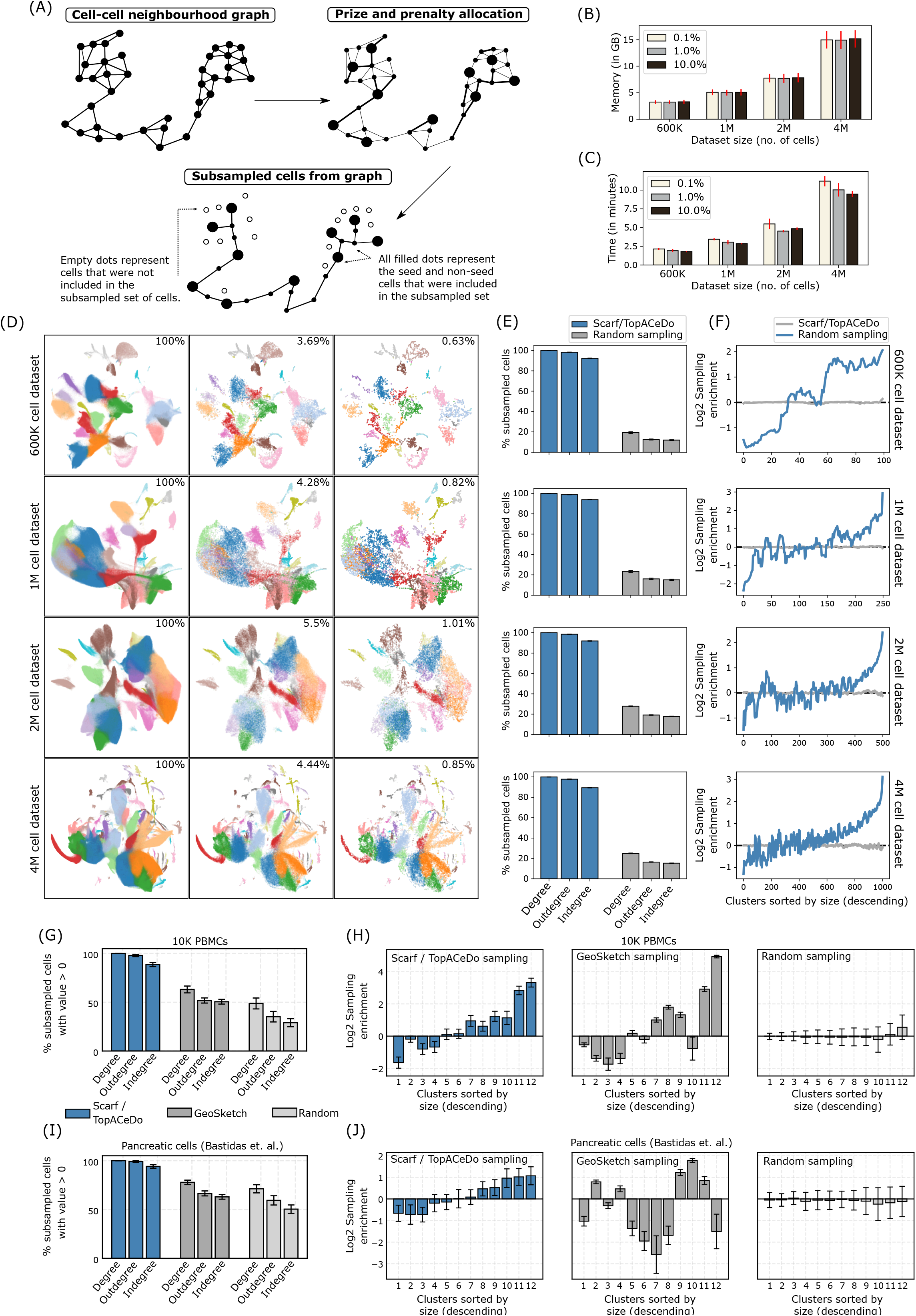
Subsampling cells from large datasets using Scarf. **(A)** An illustration of the concept behind the downsampling performed using Scarf. Each subplot represents the cell-cell neighbourhood graph computed using Scarf. The nodes in each graph indicate the cells in the dataset. The edges connect two cells, if at least one of them is present in the other’s k nearest neighbour list. **(B)** Memory and **(C)** time consumption of TopACeDo in four atlas scale datasets. The x-axis represents the number of cells in each dataset. The error bars indicate standard deviation computed using 10 iterations of downsampling. The on-figure legends show that downsampling was performed with maximum sampling rate set at 0.1%, 1% and 10%. **(D)** Scatter plots showing UMAP embedding of the four atlas scaled datasets. For each of the datasets, UMAPs with all the cells, or cells downsampled with maximum sampling rate of 10% and 1% are shown. The effective sampling percentage is indicated on each subplot. The cell colours indicate the cluster identities of cells obtained using Leiden clustering. **(E)** Percentage of subsampled cells that have non-zero graph degrees. Max sampling rate for TopACeDo set at 1% and the same number of cells downsampled with TopACeDo was used for random sampling. Bars represent mean and error bars indicate standard deviation obtained using 10 runs of subsampling. **(F)** Plots (a subplot for each of the four datasets) showing change in cluster enrichment, i.e., change in proportion of number of cells from each cluster before and after clustering. The y-axis is Log2 cluster enrichment, values below 0 indicate that proportion of cells from a cluster decreased (depleted) post subsampling and those above 0 indicate that the proportion of cells from clusters have increased (enriched). The values for each cluster represent mean of 10 iterations. The numbers on the x-axis indicate cluster ID and clusters are ordered in decreasing order by size. **(G)** Percentage of subsampled cells from 10K PBMC dataset and **(I)** differentiating pancreatic cells that have non-zero graph degrees. Max sampling rate for TopACeDo set at 1% and the same number of cells as downsampled with TopACeDo was used for random sampling and GeoSketch. Bars represent mean and error bars indicate standard deviation obtained using 10 runs of subsampling. **(H)** Bar plots indicating Log2 cluster enrichment for 10K PBMC dataset and **(J)** differentiating pancreatic cells obtained using TopACeDo, GeoSketch and random sampling. Bars represent mean and error bars indicate standard deviation obtained using 10 runs of subsampling. The cluster IDs (ordered by decreasing size) are indicated on the x-axis of each subplot.

Importantly, we found that Scarf with embedded TopACeDo was able to perform downsampling on datasets with up to 4 million cells using less than 16GB of RAM (figure 2B). In addition, downsampling took less than 3 minutes on the 1 million cell dataset and less than 15 minutes in the case of the 4 million cell dataset (figure 2C). UMAP visualization of four atlas scale datasets indicates that even with downsampling to ~1% of cells, cells belonging to all clusters across the UMAP space were sampled (figure 2D).

Furthermore, for quantitative analysis of downsampling, we analysed the degree of connections subsampled cells make with other subsampled cells in the original neighbourhood graph. A high frequency of zero-degree values indicates that many cells are disconnected from other subsampled cells and is a marker of poor subsampling, indicating that intermediary cell states are missing in the subsampled set. When comparing the number of disconnected cells between Scarf with randomly subsampled cells from four atlas-scale datasets, 100% of Scarf-subsampled cells displayed non-zero degree values across all datasets, while random sampling resulted in non-zero degree values in 19.6% to 27.4% of cells. (figure 2E).

The primary objective of downsampling is to decrease redundancy in the dataset while preserving the rare cell types/states. These two objectives can be accessed by calculating the change in the proportion of cells from each cluster after downsampling. We show that across the four atlas scale datasets, Scarf was able to reduce the proportion of cells from larger clusters while simultaneously increasing the proportion of cells from smaller clusters (figure 2F). The proportion of cells from the smallest cluster in each of the datasets increased between 5.55 and 40.48 folds while the proportion of cells from the largest cluster decreased between 2.45 and 5.12 folds. In comparison, random clustering did not show any increase in the proportion of smaller clusters or decrease in the proportion of larger clusters. As a result, random clustering has a low probability to sample rare clusters. For example, the smallest cluster in the atlas-scale datasets had none of the cells sampled in 60% (1M cells dataset), 60% (2M cells dataset), and 40% (4M cells dataset) (n=10) of random samplings. In contrast, all clusters were sampled using Scarf, regardless of the original cluster size.

In order to compare the results of downsampling between Scarf and GeoSketch, we used two contrasting small scale datasets consisting of either 10K PBMC cells of distinct cell types (provided by 10X Genomics) or 3.5K pancreatic cells ^26^ within a continuum of differentiation. Visualization of progressively increasing level of downsampling on each of these datasets, showed that Scarf was able to capture the cells throughout the UMAP landscape (supplementary figure S2). Running 100 iterations of downsampling with either Scarf, GeoSketch or random sampling, 100% of downsampled cells selected by Scarf had a non-zero degree on both the datasets, while the same measurement for GeoSketch was 63.13% and 77.79%, and for random sampling 48.77% and 71.23% in the PBMC and pancreatic cell datasets, respectively (figure 2G and 2I). Compared to random sampling, both Scarf and GeoSketch sampled an increased proportion of cells from smaller clusters and a reduction in the proportion of cells from larger clusters (figure 2H and 2J). Sampling enrichment obtained using Scarf were strongly correlated with decreasing cluster size in case both, pancreatic cells (Pearson’s r=0.98, p-value=1.66e-0.8) and PBMC dataset (Pearson’s r=0.93, p-value=7.04e-0.6), while in the case of GeoSketch, there was either no correlation (pancreatic cell dataset: Pearson’s r=0.08) or moderate correlation (PBMC data: Pearson’s r=0.79). This indicates that Scarf achieves a consistent reduction of redundancy and increase in representation of rare cells in the downsampled dataset.

### Memory efficient projection of cells for compositional analysis and co-embedding of cells

The continuous growth of publicly available single-cell genomic datasets prompts a need for reliable and appropriate data integration. In Scarf, integration takes a reference-based data alignment approach ^27–29^ wherein the transcriptional states of query/target cells are interpreted considering the heterogeneity of the reference cells. To achieve reference-based mapping, Scarf implements a highly memory efficient KNN mapping approach (see Methods). Scarf does not attempt to perform batch correction like other data integration methods ^30–32^ but rather brings the target cells into the estimated manifold of the reference cells.

Demonstrating the efficiency and accuracy of Scarf’s data integration approach, we performed mapping of published single-cell RNA-Seq dataset of interferon beta (IFN-β) treated peripheral blood mononuclear cells (n=10,111) to their culture-matched control cells (n=8,487)^33^. Prior to mapping, the datasets were independently analysed to generate UMAP embeddings of both treatment arms (figure 3B-C) and cluster annotation was performed using known marker genes of cell types previously reported for this dataset ^31^. The annotation was additionally confirmed by performing a marker gene search (supplementary figure S3A-B).

**Figure 3:**
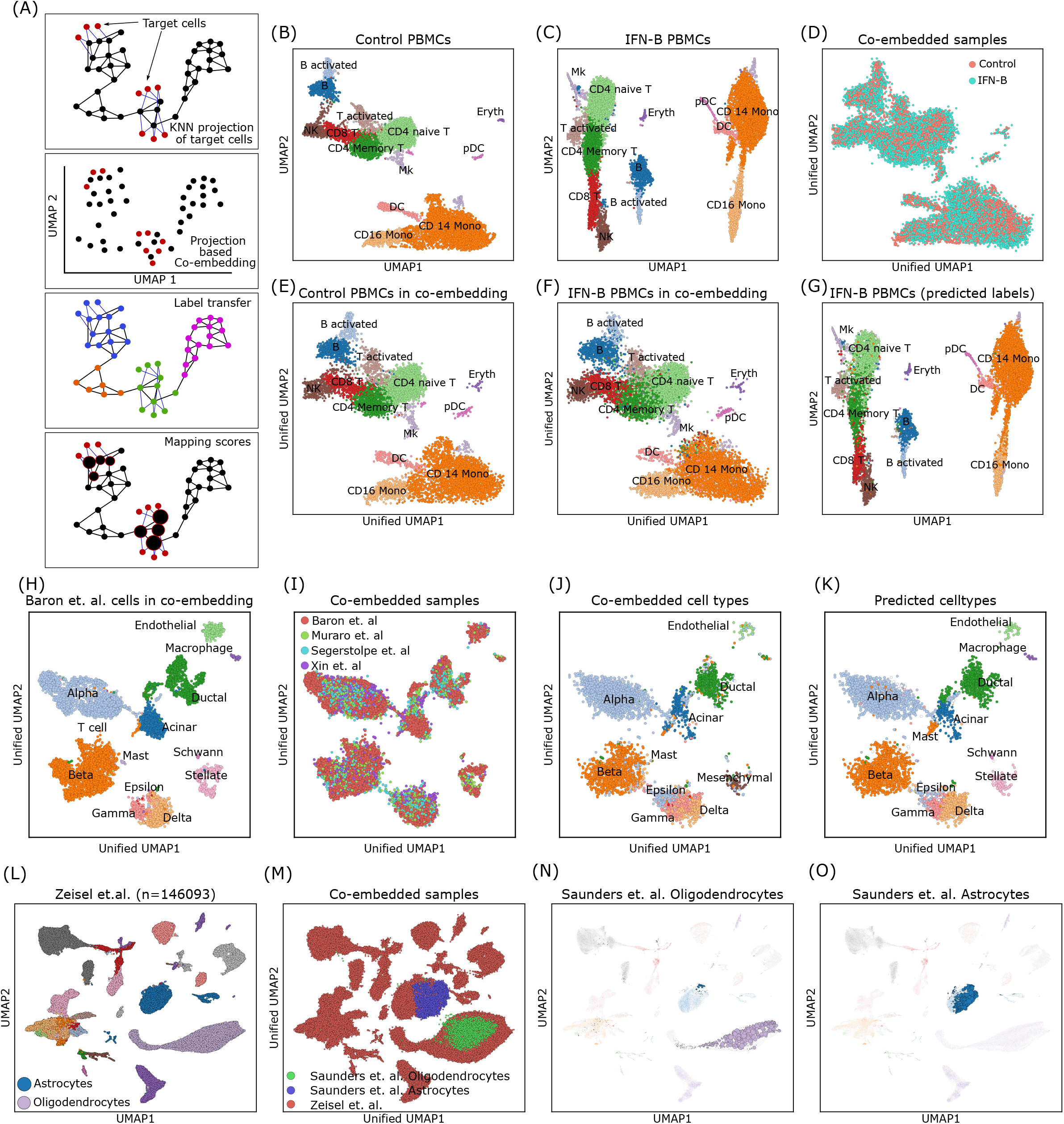
Mapping and co-embedding of cells across datasets. **(A)** The illustration shows how Scarf uses KNN projection method to co-embed projected cells into a UMAP space, then, transfer labels to projected cells and assign mapping scores to the reference cells. **(B)** Scatter plot showing UMAP embedding of untreated and **(C)** IFN-beta treated PBMCs with cells coloured based on their cluster identity; inferred cell types of each cluster are indicated **(D)** UMAP co-embedding of IFN-beta treated PBMCs post projection onto untreated PBMCs. Cells are coloured based on treatment condition. **(E)** Plot showing inferred cell types of control PBMCs in the co-embedding obtained upon projection of IFN-B treated PBMCs. **(F)** Plot showing inferred cell types of IFN-beta treated PBMCs in the co-embedding obtained upon projection onto control PBMCs. **(G)** UMAP embedding of IFN-beta PBMCs only; cells are coloured to indicate the cell types obtained through label transfer from control PBMCs upon projection. **(H)** Scatter plot showing pancreatic cells from Baron et. al. in the co-embedding space that was created using the same cells as reference. The projected cells are not shown here, and the reference cells have been coloured according to author annotated cell types. **(I)** Co-embedding of cells after projection of cells from three datasets over cells from Baron et. al. **(J)** Cells from the three projected datasets in the unified UMAP space. Cells are coloured based on author-annotated cell types. **(K)** Cells from the three projected datasets coloured based on label transfer from the reference cells (Baron et. al.) upon projection. **(L)** UMAP embedding of cells from murine nervous system atlas (Zeisel et. al.), cells are coloured based on author annotated cell types. **(M)** Oligodendrocytes and astrocytes from Saunders et. al. co-embedded with cells from Zeisel et. al. (N-O) UMAP plots showing cells from Zeisel et. al. with the size of cells scaled to show mapping score obtained on projection of **(O)** oligodendrocytes **(N)** and astrocyte from Saunders et. al.

Next, the IFN-β cells were mapped onto the control cells by including the CORAL ^34^ correction step (see Methods) and the reference cell-cell neighbourhood graph, now spiked with target cells, was subjected to the UMAP algorithm to obtain a low-dimensional co-embedding including both data sets (see Methods). The ‘unified’ UMAP of the control and IFN-β treated PBMCs (figure 3D) shows that the cells from the two datasets are well integrated. We noted that the reference cells in the unified UMAP (figure 3E) had a very similar layout as compared to the UMAP resulting from the reference cells alone. This indicates that the inclusion of the IFN-β treated cells did not disrupt the manifold of the control cells. Visualization of the cell type identity of the mapped IFN-β cells on the unified UMAP clearly showed that Scarf was able to co-embed cell types from the two datasets accurately (figure 3F).

Scarf uses the KNN mapping to transfer labels from reference to target cells (see Methods). The original UMAP of IFN-β cells (figure 3G) annotated with predicted cell types clearly shows that the label transfer was highly accurate (supplementary table ST1). Of note, Scarf predicted cell type with high accuracy even on low-abundant cell populations within the dataset. For example, 93.2% and 97.6% of the predicted cell-type labels for pre-dendritic cells and erythrocytic IFN-β treated cells respectively, were true.

Reference based data integration using Scarf is not limited to two samples. Multiple samples can be co-embedded into the reference manifold simultaneously, here demonstrated using four datasets of pancreatic cells ^35–38^. These data are from four different labs, and derived using four different single-cell RNA sequencing platforms viz., InDrop^35^ (n=7715), CEL-Seq2^36^ (n=1946), SMART-Seq2^37^ (n=1809) and C1-IFC SMARTer^38^ (n=1446). Each dataset was processed individually, and the heterogeneity was explored by generating individual UMAP embeddings. (supplementary figure SC-F). Subsequently, we used author-provided cell type labels to annotate the cells. Choosing the InDrop dataset as reference, we mapped the other datasets using CORAL corrected values and generated a unified UMAP, as done above for the PBMCs. Again, the UMAP embedding of the reference cells remained largely unaltered when co-embedded with the other datasets (figure 3H) Moreover, the co-embedding showed mixing of the datasets into each of the reference clusters (figure 3I) and visualization of the cell type identity of the mapped cells on the unified UMAP revealed that co-embedding was highly cell type specific (figure 3J). As previously described for PBMCs, the mapped cells were accurately classified into reference cell type labels (figure 3K, supplementary table ST2). For example, a rare population of endothelial cells from CEL-Seq2 (n=19) and SMART-Seq2 (n=15) datasets was labelled to 100% accuracy by the classifier while alpha cells, the most abundant cell type, were identified with approx. 99% accuracy in each of the three mapped datasets.

Having assessed the accuracy of the data integration, we aimed to assess the scalability of Scarf to larger datasets. Here, we chose a modestly large dataset with 146,093 cells from the mouse nervous system ^39^. The dataset was processed to generate a UMAP and annotated with user provided cell type specification (figure 3L). Next, 43,954 oligodendrocytes and 21,885 astrocytes from another brain cell atlas ^40^ were mapped to this reference dataset. By utilizing the nearest neighbour index that was already generated for the reference dataset, the mapping, for both the datasets took approx. 40 seconds and was done under 500MB of memory consumption. Convincingly, the unified UMAP showed that the mapped cell types colocalize with the equivalent clusters in the reference dataset (figure 3M). KNN classification was able to accurately predict the correct cell type for 97.6% of astrocytes and 99.5% of oligodendrocytes. Generating co-embedding of large datasets can be time consuming when multiple mappings on multiple large datasets are performed. To this end, Scarf provides an additional alternative in the form of mapping scores^41^ (see Methods). Here, we demonstrate individually generated mappings for astrocytes and oligodendrocytes, visualized by increasing cell size in proportion to their mapping score (figure 3N-O). Mapping score clearly indicate the reference cells that bear highest similarity to the projected cells. It was found that in the case of projected astrocytes generated 94.05% of all mapping in the astrocyte cluster of the reference dataset. Similarly, oligodendrocyte projection generated 99.49% of all mapping scores in the oligodendrocyte containing cluster of the reference database.

Taken together, Scarf presents multiple methods to perform reference-based integration in highly memory efficient way. Scarf allows caching and reusage of training indices for KNN to allow rapid mapping of datasets in cold start scenarios. Using label transfer, co-embedding and mapping scores users can quickly identify relationships between cells across atlas scale datasets.

### Memory efficient hierarchical clustering and t-stochastic neighbour embedding for single-cell genomics data

Visualization of single-cells on two/three-dimensional space is one of the central tenets of single-cell data analysis. Since single-cell datasets contain tens to hundreds of thousands of features, non-linear dimension reduction algorithms have been considered as ideal solutions for 2D/3D data visualization. Historically, the most popular choice for a variety of single-cell datasets has been t-SNE ^42^, while UMAP ^14^ is the most common current method for visualization on account of its computation efficiency (runtimes) and improved preservation of global structure in the data. Alternatively, FIt-SNE ^43^ is an improved t-SNE algorithm with runtimes on par with UMAP for larger scale datasets. Recently, a novel t-SNE algorithm, SG-tSNE ^15^, that can directly operate on stochastic KNN graphs was introduced. The implementation of SG-tSNE was shown to have improved runtimes over FIt-SNE and, to be scalable to three-dimensional embedding of large-scale datasets. However, it is not known how the embeddings obtained using SG-tSNE compared to those from UMAP and how SG-tSNE performs on a wider selection of datasets.

The neighbourhood graph computed by Scarf can be directly fed into either UMAP or SG-tSNE algorithm. Furthermore, Scarf uses the same initial embedding for both UMAP and SG-tSNE (see Methods), thus restricting the differences between the two methods to the underlying algorithm itself and the parameters used. We benchmarked the run-time of UMAP and SG-tSNE on four atlas scale datasets of 600K, 1M, 2M and 4M cells at multiple iterations (see Methods for details). Both algorithms run in an iterative manner for a user defined number of steps. Figure 4A shows 2D UMAP (250 iterations) and SG-tSNE (2,500 iterations) plots obtained of the four atlas scale datasets. SG-tSNE consumed less time for 2500 iterations compared to UMAP’s 250 (7.0, 11.8, 15.9, and 29.5 mins vs. 14.9, 30.3, 37.7, and 96.2 mins, for the 600K, 1M, 2M, and 4M cell datasets respectively). Moreover, we observed that for the corresponding iterations across datasets, SG-tSNE scaled better to the increasing number of cells; for 4M cell datasets the runtime of 10,000 iterations SG-tSNE (96.45 minutes) was similar to the runtime of UMAP with 250 iterations (figure 4B).

**Figure 4:**
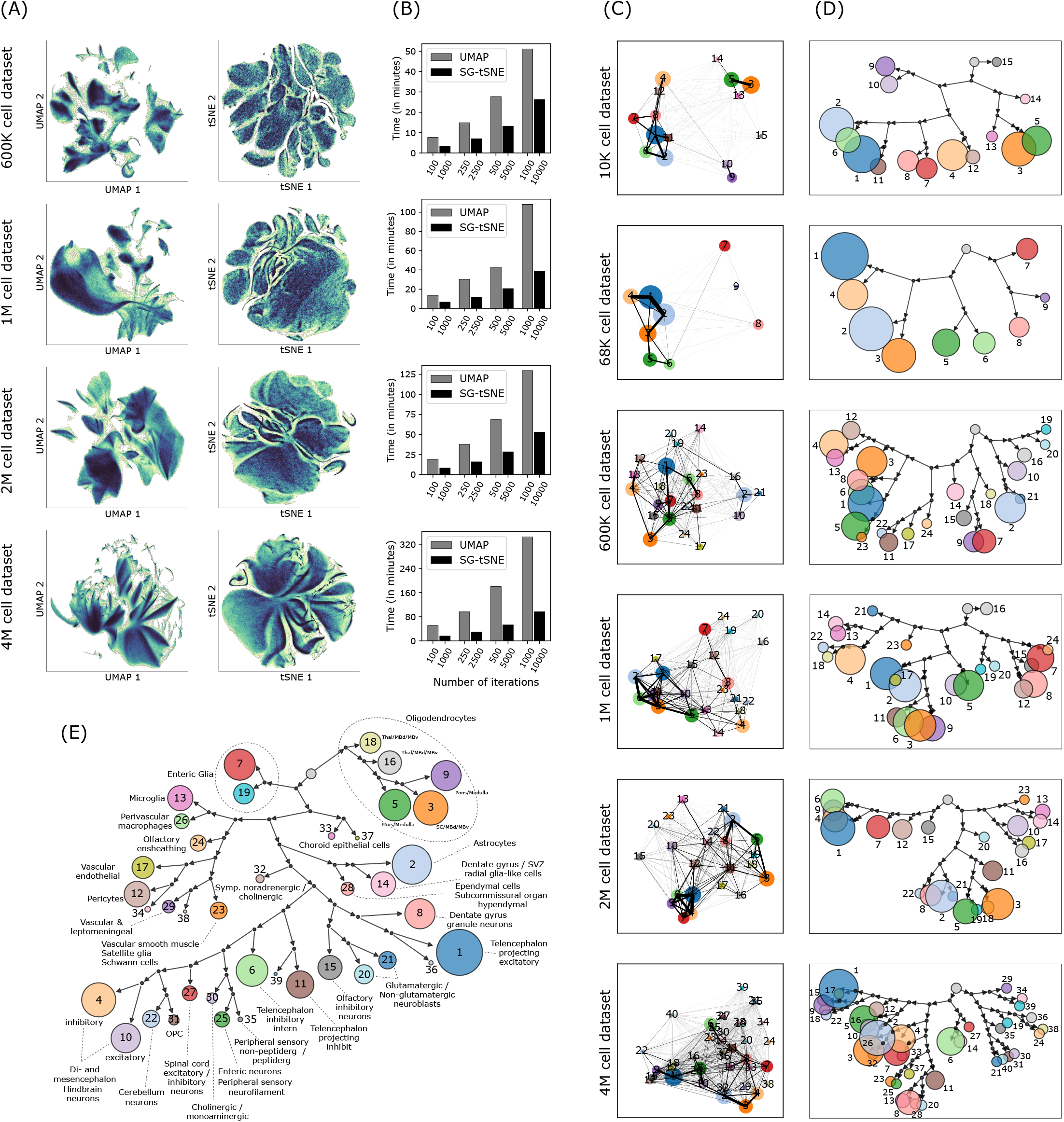
SG-tSNE and Paris hierarchical clustering on single-cell datasets. **(A)** UMAP and SG-tSNE embeddings of the four, atlas scale single-cell RNA-Seq datasets. The colour gradient shows regions of high (darker regions, with black at extreme) and low cell density (lighter regions with light yellow at extreme). **(B)** Runtime (in minutes) of UMAP and SG-tSNE at different number of iterations, on Scarf computed cell-cell neighbourhood graph. **(C)** Scatter plots, for each dataset, showing centroids of each Paris cluster in UMAP space. The lines connecting pairs of cluster centroids indicate the weighted sum of edges (in the cell-cell neighbour graph) shared by those clusters. Thicker lines indicate larger degree of similarity between the clusters. The number on cluster centroids indicate cluster ID. **(D)** Paris dendrograms of each of the datasets. Each terminal node in the dendrogram represents a cluster of cells (same cluster identities as in C above). The size of cluster node is set proportionate to the number of cells in that cluster. Each binary branchpoint in the dendrogram is shown with black circle. The root node is shown as unlabelled grey node and does not have an incoming arrow. **(E)** Paris dendrogram of the mouse nervous system cell atlas (Zeisel et al.). The coloured nodes indicate a Paris cluster and are labelled with cluster IDs. The nodes are sized proportionately to the number of cells in the cluster. The author annotated cell types present in each cluster are indicated next to the cluster node.

Annotation of UMAP (supplementary figure S4A) and SG-tSNE (supplementary figure S4B) plots with Leiden clusters resulted in well separated clustering in both cases and the relative distances between clusters could be easily visualized on the UMAP plot. Moreover, the clusters were positioned in an equivalent vicinity to other clusters in both embeddings. To quantify this, we calculated the Spearman rank correlation between the nearest neighbour across the clusters in the embedded space and the original cell-cell neighbourhood graph (supplementary figure S4C). The Spearman correlations coefficient values across the clusters show that UMAP was able to learn cluster relations quickly and showed only a slight improvement from increased number of iterations. The largest improvement was seen in the case of the 4M cell dataset where the mean Spearman’s rho value increased from 0.78 (100 iterations) to 0.84 (1,000 iterations). In all four datasets, SG-tSNE showed a smaller mean Spearman’s rho value than UMAP at lower number of iterations but improved to be larger than UMAP when higher number of iterations were used. For example, in the case of the 4M cell dataset, the SG-tSNE’s value increased from 0.63 (1,000 iterations) to 0.93 (10,000 iterations).

Next, we compared UMAP and SG-tSNE for the preservation of local data structure by quantifying the number of KNN neighbours found in the vicinity of each cell in the UMAP or SG-tSNE space (supplementary figure S4D). Across the four datasets, SG-tSNE led to improved local structure with the increasing number of iterations while no benefit was observed for UMAP embeddings. UMAP, with 1,000 iterations, had 0.14, 0.19, 0.21, and 0.24 mean fraction of nearest neighbours preserved in the embedding of 600K, 1M, 2M, and 4M cell datasets, respectively. SG-tSNE, on the other hand, at 10,000 iterations, reached the values of 0.33, 0.46, 0.44, and 0.40, in the respective datasets.

Clustering is one of the most critical aspects of single-cell data analysis wherein cells are partitioned into distinct groups. Hierarchical clustering-based methods ^44–48^ can provide accurate partitioning of cells and discover rare cell types in the data but are not scalable to atlas size datasets ^49^. Alternatively, Graph clustering ^50–53^ methods have become a popular choice for clustering on single-cell data and Louvain ^52^ and Leiden ^50^ algorithms are routinely used through Seurat ^29^ and Scanpy ^16^ packages. However, unlike hierarchical clustering methods, Louvain and Leiden are unable to clearly reflect the relationship between clusters. Scarf performs clustering using the highly scalable Leiden algorithm. Like other graph-based methods and unlike hierarchical clustering, Leiden is unable to explicitly indicate the relationships between clusters. PAGA ^54^ was introduced as a method that can explicitly indicate relationships between clusters once created by any clustering method. Unfortunately, it can become hard to interpret those relationships on larger datasets where multiple clusters with complex connection patterns are present (figure 4C). Within Scarf, we introduce a new approach, Paris ^13^, that has not previously been applied to single-cell genomics data. Paris is a graph clustering approach that rather than creating flat clusters of cells, creates a dendrogram of cells akin to hierarchical clustering methods.

Specifically, Paris computes a binary tree representing the various nested clusters of the graph, at different levels of granularity. The top-level clusters give the core structure of the graph while the lower-level clusters provide the local structure. Like Louvain and Leiden, Paris is a scalable algorithm applicable to large graphs with millions of cells. High performance is achieved through the nearest neighbour chain algorithm, where pairs of clusters to be merged are locally searched, following the edges of the graph. The successive merged clusters are stored and aggregated to compute the final binary tree.

We applied Paris to the six datasets used previously (supplementary figure S4E). In order to allow comparison with Leiden clustering (supplementary figure S4F), we cut the Paris dendrogram to obtain the same number of clusters as Leiden clustering. Coalesced Paris dendrograms presented a clear structure of cluster relationships (figure 4D) and the mean concordance between the Paris and Leiden clusters was found to be between 88.05% and 93.73% across the datasets (supplementary figure S4G). These values were significantly larger (p-value <4.25e-7; Mann Whitney U test) than randomly shuffled cluster identity wherein mean concordance with Leiden ranged between 9.92% and 26.47%.

In order to ascertain the biological meaningfulness of Paris dendrograms, we used scRNA-Seq data from cells of the murine nervous system ^39^. The authors manually curated multiple taxonomy of cell types in the cell atlas. Importantly, Paris was able to retrace the taxonomic relationships between the cell types (figure 4E) presenting a dendrogram with relevant and clearly separated branches for vascular, immune, glial, and neuronal cell types.

Together, these results show that Scarf can perform all critical analysis steps of single-cell genomics data including filtering, normalization, feature selection, linear dimension reduction, neighbourhood graph creation, embedding using UMAP and t-SNE, clustering, downsampling, and cell projection on even the largest available data sets on a regular laptop in a time efficient manner. Importantly, the resulting data are consistent or improved compared to using memory-exhausting current state-of-the-art tools.

## DISCUSSION

The ability to handle and process large-scale single-cell genomics data with readily available hardware is urgently needed. Most atlas-scale projects resort to providing online interfaces with UMAP/t-SNE plots over which users visualize the expression of different genes. But this can be grossly insufficient for many research tasks. Researchers often generate datasets that need co-analysis with other atlas-scale datasets which necessitates the use of high-performance computing machines. Another common-usage scenario where researchers need to process count matrices of large-scale datasets is when performing custom sub-selection of cells and generating new UMAP/t-SNE and clustering. Feature selection on a sub-selection of cells can improve the resolution of UMAP/t-SNE and increase the sensitivity of rare cell-type and cell-state identification. We designed Scarf to allow researchers the possibility for analysis and re-analysis of atlas-scale datasets on their laptop computers.

Coupled with memory efficient methods, like Kallisto Bustools ^55^, for generating cell-gene count matrices, Scarf represents an end-to-end solution for the analysis of single-cell RNA-Seq datasets. During the preparation of this manuscript an R-based memory efficient tool, ArchR ^56^, was published for analysis of scATAC-Seq data. In the paper describing this feature-rich tool, the authors benchmarked a simulated PBMC dataset with 1.2 million cells and found the memory usage to be over 20GB in a small infrastructure setting. With Scarf, we were able to perform corresponding steps in scATAC-Seq dataset with 700K cells within 5GB RAM. Additionally, as of the writing of this manuscript, ArchR needs genome aligned fragment files and unlike Scarf, does not support direct input of precalculated cell-peak matrices that are readily available for many atlas scale datasets. Just like Scarf has an upstream dependency on the cell-peak matrix, ArchR is dependent on external tools for genome alignment, a step that might not be memory efficient or feasible on laptop computers due to long runtimes and space requirements. Hence, neither of the currently available software yet represents an end-to-end solution for scATAC-Seq analysis.

Our novel downsampling method is designed to leverage the fact that a cell-cell neighbourhood graph is already calculated in a memory efficient way. The obtained downsampled set can thereby be exported and used with external tools to apply methods that are not currently directly supported in Scarf. We allow seamless import and export of data between Scarf and Scanpy. Though our comparison metrics indicate that Scarf performs better than GeoSketch, we want to highlight that GeoSketch does not require clustering information that Scarf/TopACeDo needs to perform downsampling.

As previously described ^57^, the embedding obtained from t-SNE and UMAP can be sensitive to the parameters used in the analysis. Hence, one method is not necessarily better than the other, but rather complementary techniques for visualization. Without extensive parameter tuning, SG-tSNE can generally reveal the heterogeneity of large-scale datasets more clearly than UMAP, while on the other hand UMAP can be more useful to investigate the relationship between clusters and visualize biological processes like differentiation. Our contribution here was to, for the first time, perform a head-to-head comparison of UMAP and a graph-based t-SNE (SG-tSNE) on multiple datasets by providing the exact same KNN graph and coordinate initialization.

In Scarf, we introduce a generic graph-based hierarchical clustering method that can clearly highlight cluster similarity. Single-cell specific hierarchical clustering algorithms have been shown to not be scalable to large-scale datasets ^48^. Our benchmarks showed that Paris algorithm applied on Scarf computed cell-cell neighbourhood graph creates a full dendrogram of a 4M cell dataset in 58 minutes. (supplementary figure S4H-I). This runtime is much slower compared to Leiden clustering which generated a fixed resolution in less than 6 minutes. However, the two runtimes cannot directly be compared as Paris reveals a full profile of cell-cell relationships while Leiden provides only a static partitioning using an arbitrary ‘resolution’ parameter. In practice, analysts often run the Leiden algorithm with many different values for the ‘resolution’ parameter to ascertain an optimal number of clusters. In contrast, the creation of a cell dendrogram in Paris is parameter free and can subsequently be cut into any desired number of clusters within a few seconds. Hence, the effective runtime of the Leiden algorithm can be longer than Paris, especially on large and complex datasets.

Taken together, Scarf represents a toolkit for comprehensive and facilitated analysis and integration of even the largest single-cell datasets on desktop computers. Thus, Scarf makes single-cell genomics available to a significantly larger segment of the research community, thereby further accelerating the current single-cell era into its second decade.

## METHODS

### Design principles

#### Data storage that supports parallel processing

Scarf, though primarily intended for memory efficient analysis, provides enhanced parallel processing capabilities for single-cell genomics analysis. We chose the Zarr file format ^17^ for on-disk representation and storage of data. The benefit of using Zarr over HDF5 (used by Loom and Anndata) is its ability to perform concurrent read/write operations on the data. We use Dask ^58^ library to load and perform efficient concurrent operations on the Zarr backed data matrix. Using Blosc library we were able to efficiently compress the data matrix while still in dense format. Storing and loading compressed, chunked dense matrix allowed us to avoid sparse/dense interconversions, a strategy used by other packages to manage memory ^16,31^. The size of the chunks of count matrix saved by Zarr represents the trade-off between speed and memory-efficiency. Larger chunks allow faster processing by reducing loading cycles but have larger memory footprint than the smaller chunks. Individual chunks are processed and fed into downstream algorithms in parallel.

#### Efficient usage of CPU resources

We observed that in the Python ecosystem many libraries aggressively use all the available CPU cores. Though this is usually not an issue when running analysis on servers, it can seriously impede user’s ability to simultaneously use their laptops/desktops if all CPU cores are being used. Hence, we have extensively placed CPU usage restriction throughout the code so that only a user-determined number of CPU cores are engaged into the analysis. We have further enabled support for running non-linear dimension reduction methods, which can often be longest running step in the pipeline, on multiple CPU cores.

#### Object oriented approach

The Scarf code was written to facilitate an object-oriented programming experience for the users. As a result, the user needs to interact with only one variable for each dataset. The data and all the analysis functions are packaged into the same variable. This allows users to easily introspect the available functions and assays simply using auto completion features of Python. We believe that this strategy can help minimize unintended mistakes by users, some of whom might be very new to programming. Our approach contrasts with Scanpy and Seurat, wherein analysis functions and data reside under separate namespace.

#### Optimizations for interactive iterative analysis

Single-cell genomics analysis has several parameterized steps and estimating the optimal parameters can often be difficult beforehand. Hence, users explore their datasets with different combination of parameters. Scarf aggressively caches all the intermediate data generated. This avoids unnecessary re-computation.

#### Multi-assay support

Multi-omics experiments like CITE-Seq are well supported by Scarf. Scarf can store assays data from each of the modalities under a separate branch of the Zarr hierarchy. The cell level attributes are stored outside the assays while feature level attributes and assay’s count matrix is stored under the assay group. Scarf’s programmatic interface mimics this hierarchy. Also, Scarf assigns one of the assays as default when the dataset is loaded, and all the functions will act on this default assay. Almost all the functions associated with ‘DataStore’ object use ‘from_assay’ parameter which can be used to assign the assay for the operation.

### Scarf core steps

#### Cell filtering

Poor quality cells can be filtered based on any cell attribute. Scarf has a ‘auto_filter_cells’ function that will model a normal distribution of a chosen cell-attribute and then remove cells with value probability below x or over 1-x, wherein x is a user selected cut-off value (default value: 0.01). Users can also apply custom filtering methods to remove or include cells into the analysis.

#### Normalization

For scRNA-Seq datasets Scarf performs library size normalization of the cells. Hence, the normalized value of a gene/feature in a cell *y_Fc_* can be calculated as

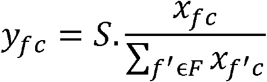

where *F* is the feature set, *c* represents a cell and *S* is a scaling factor. The normalized value can optionally be transformed into log scale: *y_fc_* = *log*(1+ *y_fc_*)

For scATAC-Seq datasets, TF-IDF (term frequency-inverse document frequency) normalization is performed.

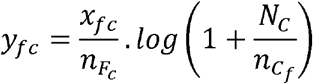

Where 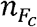 represents the total number of accessible peaks (those with non-zero values) in a given cell, 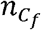 represents total number of cells where a given peak is accessible. *N_c_* is total number of cells in the dataset.

#### Feature selection

For scRNA-Seq datasets, Scarf provides highly variable gene (HVG) selection approach as previously reported^59^. For this purpose, the mean expression and variance of genes are calculated across all the cells and converted into log scale. The next step is to remove the mean-variance trend in the feature space. For this, genes are divided into equal sized bins based on their mean expression value; from each bin, the gene with lowest expression value is selected. A lowess curve (using Python’s statsmodels package ^60^) is fitted to the selected genes. The fitted curve is used to predict the ‘expected variance’ for each gene using mean expression as independent variable. The ‘expected variance’ is subtracted from observed variance to obtain corrected variance based on which HVGs are selected. Optionally, the users can put constraints on mean expression and corrected variance to perform HVG selection. For scATAC-Seq datasets, each peak is assigned a prevalence score. The prevalence score is the sum of TF-IDF normalized values for all the features. Top n peaks, sorted by prevalence scores are chosen by users to perform the downstream analysis.

#### Primary dimension reduction

For single-cell RNA-Seq datasets, PCA is applied to perform linear dimension reduction. PCA is normally applied to a subset of features prioritized using a feature selection method like highly variable gene selection but can also be applied to all the features or user designated custom subset of features. In Scarf, Sklearn’s incremental PCA implementation ^61^ is used which allows PCA to be trained iteratively using only a subset of cells at a time. Additionally, using Scarf, users can easily use only a subset of cells to train PCA and the rest of cells can still be transformed into this trained PCA space. The data is standard scaled before performing the PCA. For scATAC-Seq datasets, we use Gensim’s iterative latent sematic indexing (LSI) ^62^ method to perform dimension reduction. Similar to PCA, the LSI method can be trained iteratively and hence is scalable to extremely large datasets.

#### Cell-cell neighbourhood graph creation

Approximate nearest neighbour algorithm, hierarchical small navigable world (HNSW), as implemented in HNSWlib library is utilized ^63^. The dimension reduced values are used to build the HNSW index. By default, Scarf uses Euclidean metric as the measure of distances between the cells. The index once trained is saved onto the disk for later use. The nearest neighbour search is then performed using the index, querying for ‘k’ neighbours. During the query, it is noted if cells are indeed the nearest neighbours of themselves and this is summarized and reported as the recall value. The resulting KNN index and the distances to the KNNs is saved onto the disk. Thereafter, an edge normalization step is performed on the KNN graph wherein the distances are converted into a continuous scale bounded between 0 and 1 using UMAP’s edge weight smoothening algorithm ^64^. The resulting adjacency matrix is saved in sparse format on disk. This adjacency matrix represents as the graph that is directly fed downstream steps like, UMAP, SG-tSNE, Paris and Leiden clustering.

#### Calculation of initial embedding

Before running UMAP and tSNE, cells are usually ascribed an initial embedding which can be either random coordinates (default for many t-SNE algorithms) or informed measures such as first to/three principal component ^57^ or Laplacian eigenmaps (default for UMAP). In Scarf, the initial embedding for UMAP and tSNE is calculated by fitting MiniBatch Kmeans algorithm from scikit-learn package^61^ during the same step as the graph construction. PCA is performed on the Kmeans clusters centroid matrix which of form *n* × *c*, where *n* is the number of cells and *c* is the number of centroids.

### Benchmarking setup

All benchmarks were performed on Lunarc computing cluster comprised of nodes with configuration: 16 2.6GHz CPUs and 190 GB RAM. Each compute node had a local SSD drive and files were moved to local storage to allow exclusive IO operations for each job. A job comprised of either the Scarf (version 0.5.9) or Scanpy (version 1.6.1) pipeline which was monitored for memory consumption using Linux’s ‘ps’ utility. For benchmarking of cell downsampling, version 1.2 of GeoSketch was used.

### Marker feature identification

In Scarf, we adopt a simple and fast approach to identify marker genes based on gene ranks. All the features are ranked based on normalized values. Thereafter, for each gene, we calculate the mean rank for each group/cluster. The mean ranks for each gene-cluster are normalized by dividing by sum of all mean ranks from all the clusters. The sorted list of normalized mean-ranks gives the ordered list of marker features for each cluster.

### TopACeDO algorithm

The first step in the TopACeDO algorithm is to identify ‘seed’ cells in the graph. For this, two metrics for each individual cell (referred to as node in the neighbourhood graph) are calculated: n-neighbourhood degree (NND) and neighbourhood-connectedness (NC). The degree of the node is calculated as the total number of other nodes this particular node is connected to. 1-neighbourhood degree is the sum of degree of all nodes that are connected to a given node. Hence, NND is computed by iterating neighbours of neighbours over n-step distance and captures the density of the connections around a given node in the graph. The second metric is neighbourhood connectedness that captures if a given number of connections are shared between many or few nodes. To calculate NC of each cell, the sum of shared nearest neighbour distances (Jaccard distance) between a node and all its neighbours is calculated. Thus, if a node is connected to other nodes that are strongly connected with each other, this node will get a high value for neighbourhood-connectedness.

For the next step, the algorithm uses partitioning of cells. Here, median NND and NC is calculated for each cluster of cells and the median value is used to adjust the sampling rate for each cluster. A higher median NND leads to reduction in sampling rate while a higher NC leads to a reduced sampling rate and vice-versa. Based on the sampling rate, the number of cells to be sampled from each of the clusters are determined. Each cluster is then sub-clustered, wherein the number of sub-clusters are the same as the number of cells to be sampled; one cell is then sampled from each of the sub-clusters. These sampled cells are referred to as ‘seed’ cells.

All the seed cells are assigned a constant prize value. Here, we used value of 10. The edge penalty *E_p_* for each edge is calculated as following:

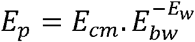

Wherein, *E_cm_* and *E_bw_* are user provided parameters, edge cost multiplier and edge bandwidth, respectively and *E_w_* is the edge weight in the graph. Higher values for *E_cm_* will make reaching remote cells in the graph more difficult but at the same time will discourage inclusion of non-seed cells in the downsampled set. Higher *E_bw_* accentuates the difference among edge penalties. Here we used *E_cm_* = 1 and *E_bw_* = 10

### KNN mapping and Integrated embedding

KNN mapping is performed using the precalculated HNSW index of the reference dataset. By avoiding recalculation of index, Scarf can quickly perform mapping even when cold started on a dataset. If the two datasets have identical populations, we suggest usage of domain shift correction method, CORAL, that’s built into Scarf. CORAL^34^ correction is performed as follows:

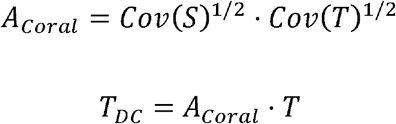

Wherein, *S* and *T* are scaled and normalized count matrixes of source and target samples respectively and *T_DC_* is the domain corrected matrix of the target dataset. Because the covariance, *Cov*, of matrices is calculated for features this normalization is easily scalable to large datasets.

All reference graphs were constructed with 11 neighbours and using 25 PCA dimensions (calculated on 2500 HVGs). The unified UMAP/SG-tSNE for the reference and target cells are calculated by spiking the reference cell-cell neighbourhood graph with target cells based on their nearest neighbours (top 3 for all the results here) in the reference set. The target-reference edges are assigned a constant weight which users can tune manually. We used a value for 1 for all the results presented. The UMAP was run for 100 iterations on the unified graph of reference and target cells to obtain a unified UMAP embedding.

To calculate the mapping scores of reference cells, we first calculate the edge weights, *W*, between reference and target cells as follows:

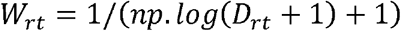

Wherein *D_rt_* is the Euclidean distance a reference and target cell. For each reference cell, mapping score, *M_r_* is calculated is as follows:

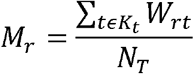

Wherein *K_t_* are all the targets that mapped to the given reference cell and *N_T_* is the number of target cells. Normalizing with *N_T_* allows improved comparison of mapping scores from two different mappings. For the reported results, the mapping scores were additionally multiplied by a scalar value of 1000 and log transformed.

Label transfer is performed in Scarf using the KNN projection. To ascribe a reference label to a target (projected) cell, we compute at the weighted sum of all edges from a target to all the connected reference classes. If the target cell has at least 50% (default threshold and used throughout here) of all the edge weights a single reference class then the target cell is ascribed to that reference class, otherwise it labelled as NA (null value meaning not assigned).

### Comparison of UMAP and SG-tSNE on atlas scale datasets

We chose 4 iteration sets for UMAP (100,250,500 and 1000) and ten times for SG-tSNE. We observed that individual iterations of SG-tSNE had lower runtime that UMAP, hence we chose iterations so that they can have comparable runtimes. Both the methods were run in parallel mode using 16 computing cores. Parameters used for UMAP: *min_dist=0.5, spread=2.0*. Parameters used for SG-tSNE: *alpha=50, box_h=1*. To find the KNN of cells in the UMAP/SG-tSNE space, we used NearestNeighbors function from scikit-learn package with Euclidean distance metric and *KD tree* algorithm. To calculate the Spearman’s correlation between the clusters in neighbourhood graph and UMAP/tSNE space, we took the following approach. We first created a similarity matrix of dimension (P,P) where P is the number of clusters identified in the dataset. This similarity matrix is calculated by calculating the weighted sum of edges (in cell-cell neighbourhood graph) between cells of each pair of clusters. Another similarity matrix is calculated based on the KNN graph calculated on the UMAP/tSNE embedding. Spearman’s correlation coefficient is calculated between each column (cluster) from the two similarity after log2 transforming the values.

## DATA AND CODE AVAILABILITY

The source code for Scarf package is available here: github.com/parashardhapola/scarf. The documentation for installation and usage of Scarf can be found here: scarf.readthedocs.io. Download links for all count matrices can be obtained using following command: ‘*from scarf.bio_data import datasets*’. The count matrices can be downloaded using command: ‘*scarf.fetch_dataset(sample)*’. Where sample is unique id of dataset in *datasets*

---

**Figure S1:**
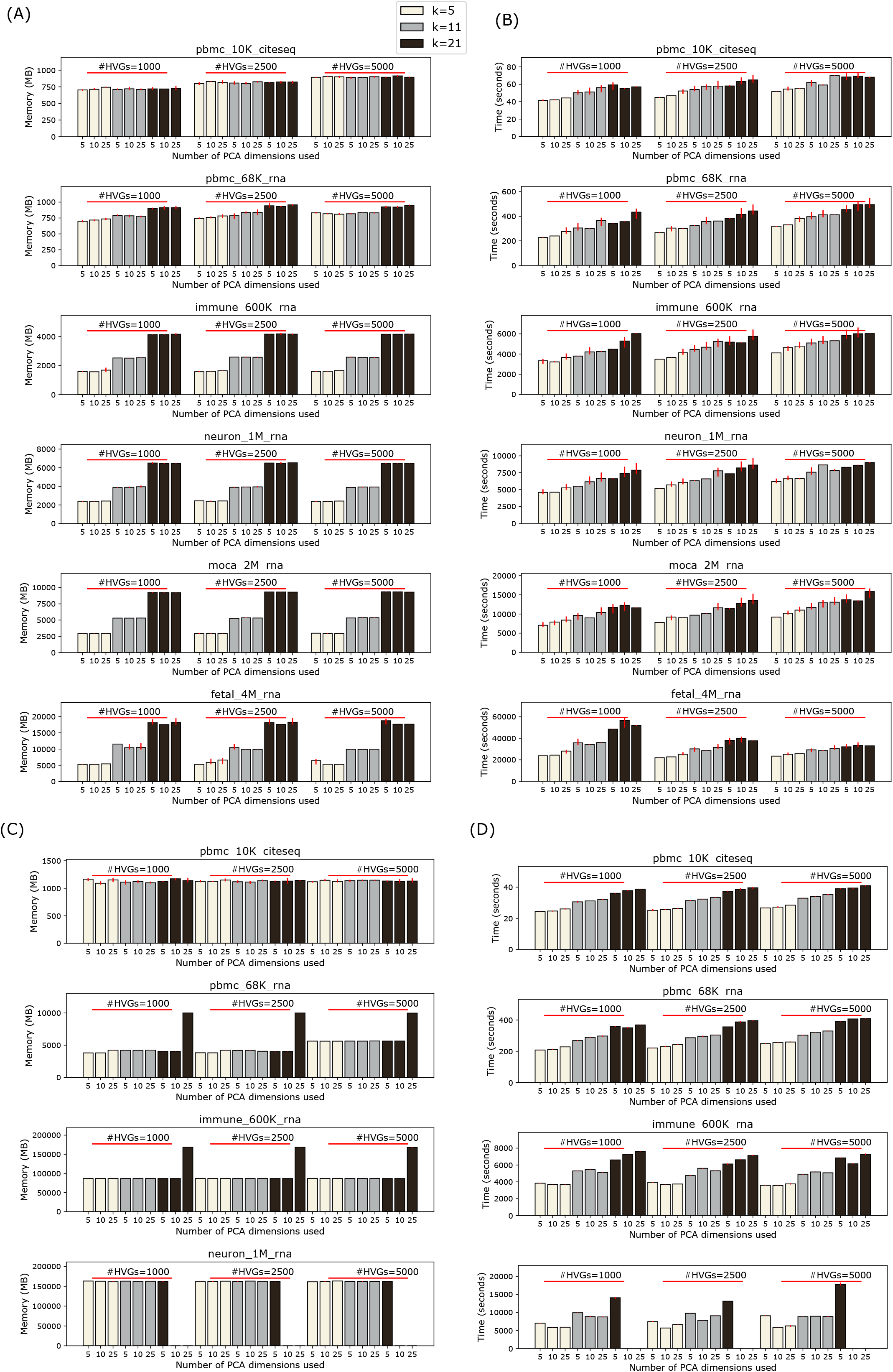
Benchmarking time and memory usage of Scarf and Scanpy across different parameters. Barplots showing **(A)** time and **(B)** memory usage of Scarf and **(C)** time and **(D)** memory usage of Scanpy across different datasets used in the analysis. Error bars show standard deviation calculated using three replicates. Each bar indicates time or memory usage under combination of number of HVGs used to perform PCA, value of parameter k (nearest neighbours in KNN graph) and number of PCA dimensions used to create the KNN graph.

**Figure S2:**
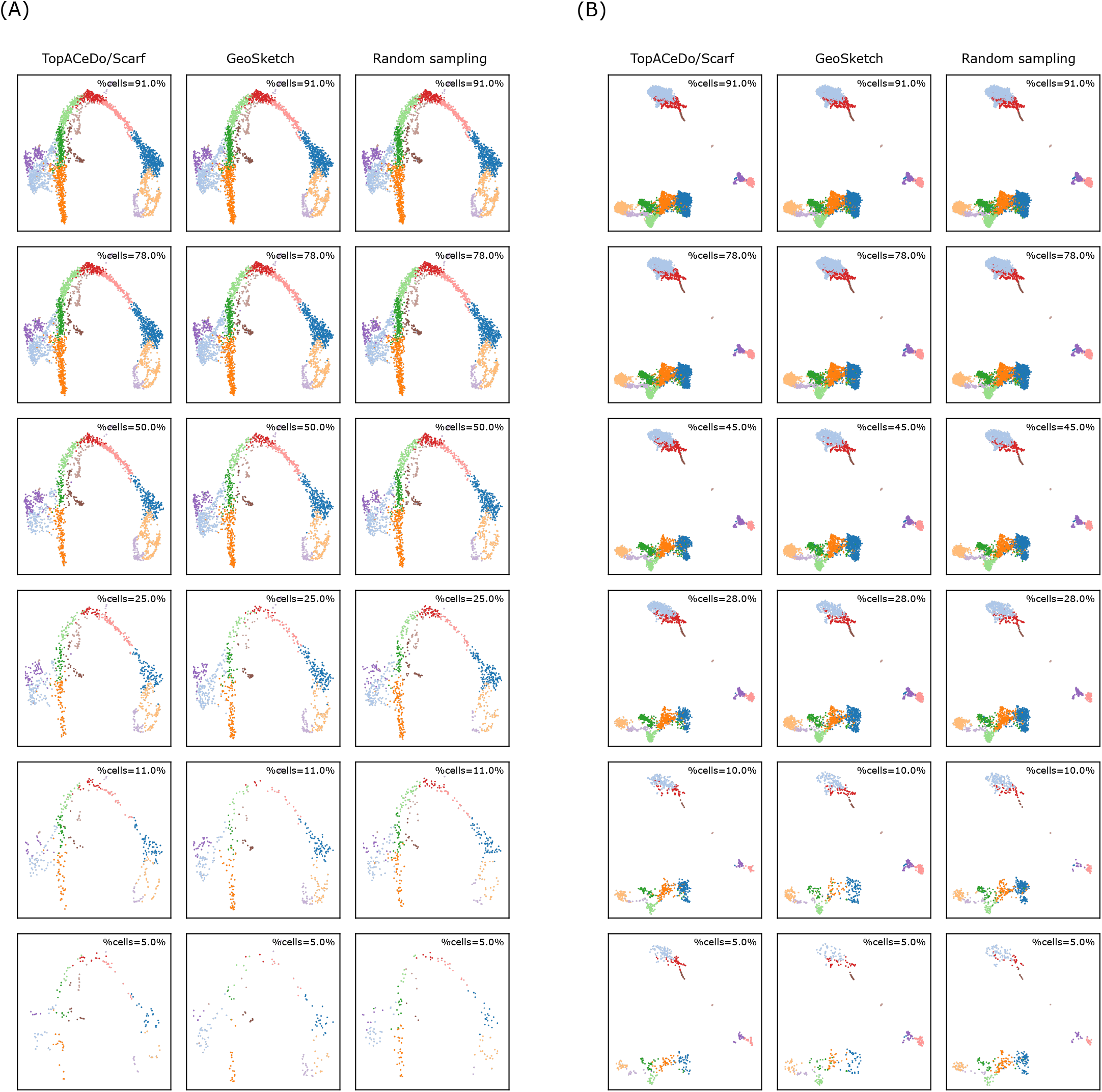
Multiple levels of down sampling using TopACeDo algorithm. **(A)** UMAP plots showing downsampled cells after applying TopACeDo, GeoSketch and random sampling to Bastidas-Ponce et. al. and **(B)** 10K PBMC dataset. The plots are arranged to show a progressively increasing degree of downsampling.

**Figure S3:**
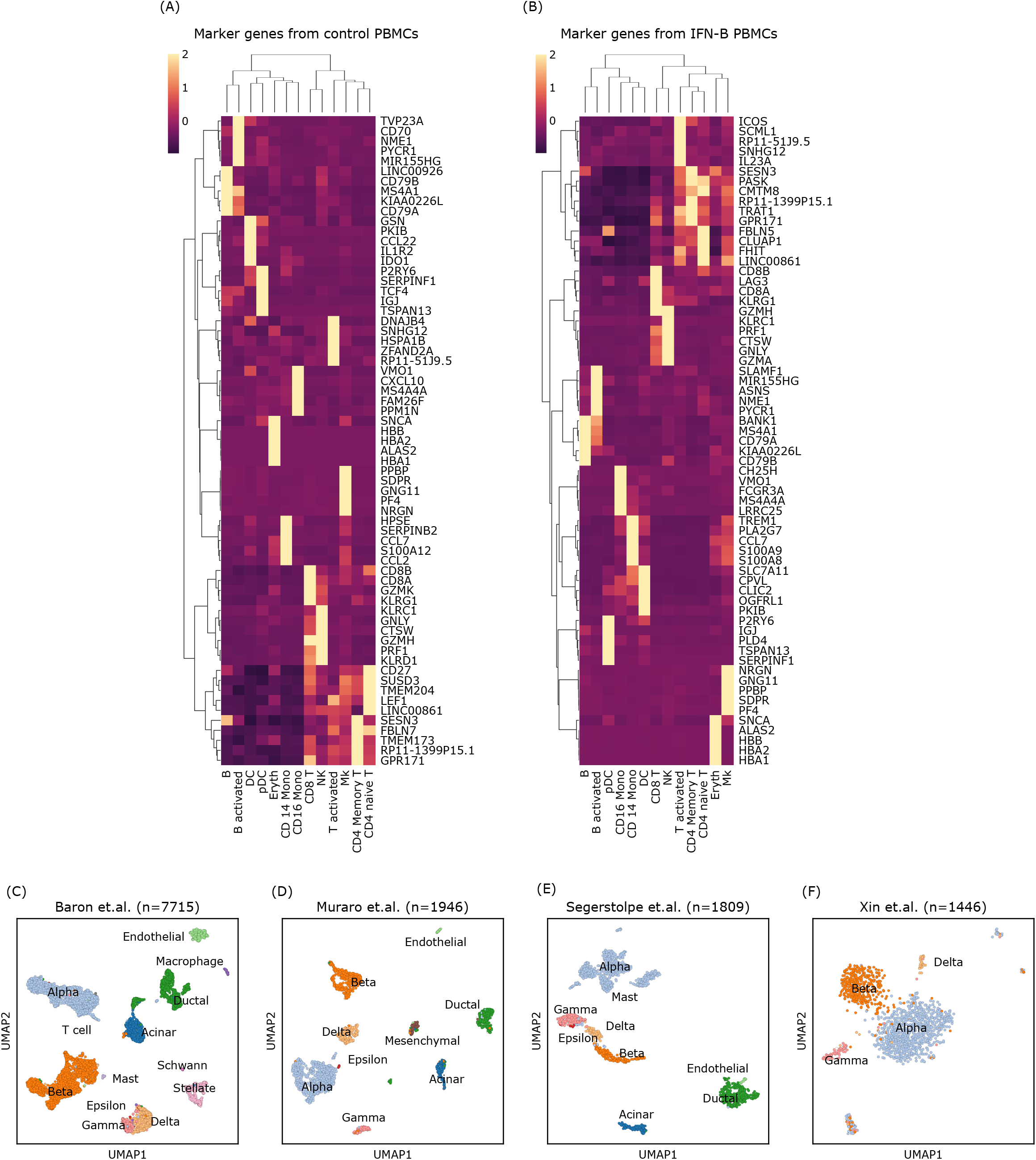
Data integration using Scarf. **(A)** Heatmap showing z-scaled normalized expression values of top five marker-genes of each cluster from untreated PBMCs from Kang et al. and from **(B)** IFN-B treated PBMCs. UMAP embedding of cells showing original author annotated cell type labels from **(C)** Baron et al. **(D)** Muraro et. al. **(E)** Segerstolpe et al. and **(F)** Xin et al. The number in the subplot title indicate the number of cells in the embedding.

**Figure S4:**
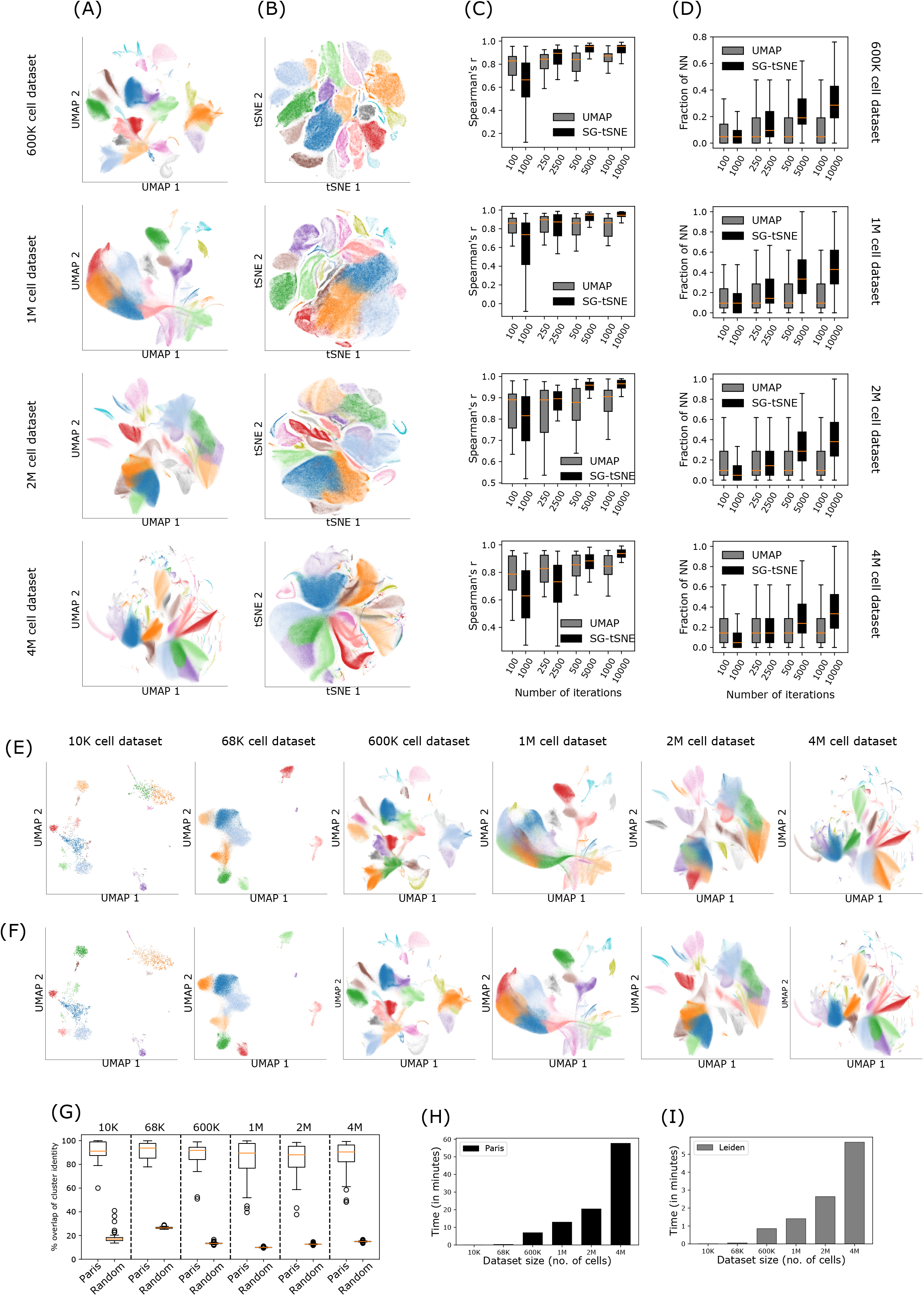
Comparison between UMAP and SG-tSNE; and Paris and Leiden clustering algorithms. **(A)** Plots showing UMAP embedding (250 iterations) of atlas scale datasets. **(B)** Plots showing SG-tSNE embedding (2500 iterations) of atlas scale datasets. For both UMAP and SG-tSNE, cells are coloured based on Leiden cluster labels. The plots show density of cells at each pixel to prevent overplotting of cells. **(C)** Boxplots showing distribution of Spearman’s coefficient values calculated between similarity matrices (cluster by cluster) of cell-cell neighbourhood graph and shown embeddings, across four atlas scale datasets. The median values are marked using red lines in the boxes. The x-axis shows progressively increasing number of iterations of UMAP and SG-tSNE that were used to generate the embedding. **(D)** Boxplots showing distribution of fraction of nearest neighbours that were conserved in the labelled embeddings, across four atlas scale datasets. The median values are marked using red lines in the boxes. **(E)** Plots showing UMAP embedding of cells, coloured according to cluster identity obtained using Paris and **(F)** Leiden clustering algorithm, across six datasets. **(G)** Boxplots showing the percentage overlap between cluster identity of cell between either Leiden and Paris or Leiden and random clustering. Each data point in the boxplot represents a Leiden cluster. The approx. number of cells in the datasets are indicated as subplot labels. **(H)** Barplots showing the time consumption (in minutes) of Paris and **(I)** Leiden clustering on the six benchmarked datasets.

## AUTHOR CONTRIBUTIONS

PD and GK conceived and designed the study; PD designed and performed all the analysis with consultation from GK.; PD, RO and JR wrote and debugged the vignettes; PD prepared the figures with consultation from GK, EE, RO and SS; GK, PD, TB and EE wrote the manuscript with assistance from SS; GK supervised the study; All authors reviewed, edited and approved the manuscript.

## ACKNOWLEDGEMENTS

This work was supported by grants from the Swedish Cancer Society, The Ragnar Söderberg Foundation, the Knut and Alice Wallenberg Foundation, the Swedish Research Council, the Swedish Society for Medical Research, and the Swedish Childhood Cancer Foundation. We thank members of Soneji lab and Stem Cells and Leukemia lab at Lund University, for their constructive suggestions throughout the development of this project.

